# Successful Demonstration of Assisted Gene Flow in the Threatened Coral *Acropora Palmata* Across Genetically-Isolated Caribbean Populations using Cryopreserved Sperm

**DOI:** 10.1101/492447

**Authors:** Mary Hagedorn, Christopher A. Page, Keri O’Neil, Daisy M. Flores, Lucas Tichy, Valérie F. Chamberland, Claire Lager, Nikolas Zuchowicz, Kathryn Lohr, Harvey Blackburn, Tali Vardi, Jennifer Moore, Tom Moore, Mark J. A. Vermeij, Kristen L. Marhaver

## Abstract

Global change will compromise the population sizes, species ranges, and survival of economically-important plants and animals, including crops, aquaculture species, and foundational ecosystem builders. Scleractinian reef-building corals are a particular concern because they are slow-growing, long-lived, environmentally-sensitive, and concentrated in the warmest regions of the ocean. Assisted Gene Flow (AGF) is considered a viable tool to help natural plant and animal populations, including corals, adapt to changing environments. Our goal was to test for the first time whether cryopreserved coral sperm could be used to facilitate assisted gene flow between genetically-isolated populations of a Caribbean coral. We collected, pooled, and cryopreserved coral sperm from the threatened Caribbean coral *Acropora palmata* in the western Caribbean (Key Largo, FL), central Caribbean (Rincón, Puerto Rico), and eastern Caribbean (Curaçao). Alongside freshly-collected sperm from Curaçao, the cryopreserved sperm from each of these populations was used for *in vitro* fertilization experiments with freshly-collected eggs from Curaçao. Across five egg donors, average fertilization success was 91 to 99% for CUR × CUR (fresh sperm) crosses, 37 to 82% for CUR × CUR (frozen sperm) crosses, 3 to 19% for CUR × FL (frozen sperm) crosses and 0 to 24% for CUR × PR (frozen sperm) crosses. Notably, fertilization was achieved in all four categories of crosses, showing for the first time through direct evidence that populations of *A. palmata* are reproductively compatible, and that genetic diversity can be transferred from one population to another for the purposes of assisted gene flow. The resulting larvae were reared in Curaçao for up to 7 days, then the swimming larvae were transported to Florida for settlement and grow-out at two separate facilities, which achieved larval settlement rates of 37 to 60% across all cohorts. Larvae were reared and settled in Florida to acclimate them to the ambient water quality, microbial environment, and temperature regimes of the western genetic *A. palmata* population as early in their life cycle as possible. At one month, over 54% all settlers had survived, including over 3500 settlers from CUR x CUR (frozen sperm), 1200 settlers from CUR × FL (frozen sperm), and 230 settlers from CUR × PR (frozen sperm). These experiments represent the first-ever pan-Caribbean coral crosses produced in captivity and the first direct evidence that geographically-separated and genetically-isolated populations of any Caribbean coral are reproductively compatible. Moreover, with over 4700 *A. palmata* settlers produced using cryopreserved sperm, this represents the largest living wildlife population ever created from cryopreserved material. Together, these findings demonstrate that cryopreservation of coral sperm can enable efficient, large-scale assisted gene flow in corals. This form of assisted migration can not only help to preserve the population-level genetic diversity of extant coral populations but also help to increase population resilience to global change.

## Introduction

As global change threatens the diversity, population sizes, population ranges, economic value, and survival of terrestrial and marine species, Assisted Gene Flow (AGF) has become recognized as an increasingly promising option for accelerating population adaptation and buffering species against extinction (Hoegh-Guldberg et al. 2008; Aitken and Whitlock 2013; Palumbi et al. 2014; van Oppen et al. 2014; van Oppen et al. 2015; Anthony et al. 2017). As a form of assisted migration, AGF is defined as “the managed movement of individuals or gametes between populations within species ranges to mitigate local maladaptation in the short and long term” (Aitken and Whitlock 2013). Although many wildlife populations are well-adapted to the specific environments they experienced over evolutionary timescales (Hereford 2009), rapid global change is now causing genotype-environment mismatches and consequent reductions in growth, survival, and reproductive health, especially in wildlife populations that are small and genetically limited (Aitken and Whitlock 2013). This is particularly true for scleractinian reef-building corals, which already occupy less than 0.25% of all ocean habitat by area and which face rapidly-warming and acidifying conditions (Hughes et al. 2018).

Under accelerating global change, some wildlife populations will acclimate by migrating to more favorable locations. However, population migration is virtually impossible for large, long-lived, sessile adult organisms such as terrestrial plants and corals, which has led conservationists to consider the feasibility of assisted migration. As a form of assisted migration, AGF is now considered a viable option to help buffer coral populations against global change (National Academies of Sciences Engineering and Medicine 2018). AGF is a particularly attractive option for corals because it enables the conservation of population genetic diversity first and foremost, with or without deliberate selection for specific phenotypic traits. Furthermore, AGF can enable the production of new genetic combinations that are more stress-tolerant under modern climate regimes. By acting on the entire genome as a whole, AGF is akin to the cross-hybridization of different populations to create robust crop varieties, livestock lineages, and forestry species, and the genetic rescue of endangered species using individuals from closely-related species. AGF is essentially a conservation-driven version of the selective breeding process that humans have applied to plants and animals for millennia.

The first practical application of AGF in corals was conducted by Dixon and colleagues on the Great Barrier Reef (Dixon et al. 2015). In their study, the authors moved live coral colonies separated by a 5° latitudinal gradient on the Great Barrier Reef, crossed the eggs and sperm from the two locations, and then demonstrated that a subset of the resultant larvae from the AGF crosses exhibited heritable heat tolerance. This study demonstrated that coral thermal tolerance is indeed heritable, and that phenotypic traits can be transferred between coral populations that span a relatively wide geographic range.

Moving any adult animals for reproductive purposes can be costly and cause stress that leads to reproductive decline. Adult coral colonies are particularly difficult to move given their size, weight, and solid attachment to the reef benthos. Furthermore, the mucus, tissue, and skeleton of a coral colony contain diverse viruses, bacteria, fungi, parasites, and algae (Knowlton and Rohwer 2003; Wegley et al. 2004; Rosenberg et al. 2007; Yarden et al. 2007; Marhaver et al. 2008; Barott et al. 2012; Pollock et al. 2018), including potential pathogens and diverse endolithic organisms (e.g., (Harvell et al. 2007; Dinsdale et al. 2008)), which have the potential to become invasive or uncontrolled if moved. The movement and introduction of marine invasive species has been known to occur through the release of ballast water from ships (Ruiz et al. 2013; Miller and Ruiz 2014) and a similar risk must be accounted for during the movement of coral colonies (Ruiz et al. 2013; Miller and Ruiz 2014). Therefore, the most practical mechanism by which to conduct large-scale AGF for coral conservation is to move gametes, thereby moving the genetic diversity but not the adult individuals.

Unlike seeds, coral eggs and sperm remain viable for only minutes to hours after they are produced (Levitan et al. 2004; Fogarty et al. 2012), meaning that freshly-collected coral gametes cannot be transported across long distances to conduct AGF. Furthermore, most coral species release gametes on only a small number of days per year (Szmant 1986; Gleason and Hofmann 2011; Harrison 2011); this increases the number of sperm-egg interactions in the water column and helps each population achieve high fertilization rates, but this compression in time dramatically limits the ability of conservationists to work with, transport, and cross-fertilize germplasm from these animals. For each spawning coral species, there are only a few hours per year during which fresh gametes can be collected and mixed. However, cryopreservation and long-term liquid nitrogen storage of coral sperm is now possible, thanks to recent advances in the collection, analysis, freezing, storage, and thawing of this material (Hagedorn et al. 2006; Hagedorn et al. 2012a; Hagedorn et al. 2017; Viyakarn et al. 2018). Moreover, in 2018, the National Academies of Sciences, Engineering, and Medicine specifically highlighted the need for well-developed and reliable cryopreservation methods to help enhance and conserve coral reef biodiversity (National Academies of Sciences Engineering and Medicine 2018). This study specifically noted that coral germplasm banks can serve not just as archives, but as tools for actively increasing genetic diversity and adaptation rates in coral populations.

Cryopreservation has been used successfully for decades to preserve both wildlife species and their genetic diversity (Tiersch and Green 2011; Holt et al. 2014). The process of cryopreservation successfully maintains live cells and tissues at ultralow temperatures through a series of dehydration and freezing steps that extract water from cells and replace the water with cryoprotectant molecules. The cryoprotectant serves as a type of antifreeze, preventing the formation of damaging ice crystals, which can otherwise compromise cell integrity and viability during the freezing or thawing process (Tiersch and Green 2011; Holt et al. 2014). The partially-dehydrated cells then can withstand the extraordinary stress of low temperature exposure and remain alive in a state of ultra-low metabolic activity (Mazur 1970; Mazur 1997; Mazur et al. 2008). Already, cryopreserved sperm has been used to facilitate reproduction in both endangered animals and livestock. Most notably, cryopreservation has enabled the propagation of endangered species including the black-footed ferret (Howard et al. 2016), cheetah (Crosier et al. 2009), and giant panda (Spindler et al. 2004).

Cryopreservation of coral sperm and fertilization of coral eggs with thawed cryopreserved sperm have both been successfully demonstrated in the past (Hagedorn et al. 2012a; Hagedorn et al. 2012b; Hagedorn et al. 2017). Coral sperm repositories in the U.S. and Australia (Hagedorn et al. 2012b) now hold material from over 25 coral species. In 2017, using two reef-building species on the Great Barrier Reef, coral larvae were successfully produced using cryopreserved sperm and the resulting larvae had the same settlement success as conspecifics produced using fresh sperm (Hagedorn et al. 2017). This study demonstrated that cryopreserved sperm can be used successfully to propagate coral juveniles up to the settlement stage. These experiments were performed using cryopreserved sperm and fresh eggs from within the same local population. To date, no studies have tested whether cryopreserved coral sperm from one location can be used to successfully fertilize coral eggs from a second, genetically-isolated location. This knowledge is fundamental to determining whether genetically-isolated coral populations remain reproductively compatible and can therefore be interbred for conservation purposes. Conversely, if completely insurmountable reproductive barriers exist between genetically-isolated populations of coral, each population must be managed and conserved as a separate biological species, and tools such as AGF cannot be used to fortify either population with genetic diversity from the other.

The Caribbean elkhorn coral *Acropora palmata* is considered one of the most ecologically-important and ecologically-imperiled species in the Caribbean. This large, branching coral occupies shallow, high-energy reef zones where it builds habitat, buffers wave action, and protects shorelines (Shinn 1980). In the 1980s, *A. palmata* and its sibling species, *A. cervicornis*, suffered a population decline of over 95% due to a Caribbean-wide outbreak of White Band Disease (Precht et al. 2002). Decades later, these species have made only modest recoveries in a handful of locations (Idjadi et al. 2006; Lidz and Zawada 2013; Crabbe and James 2014). Both species have been further compromised by habitat loss, poor water quality, physical impacts, algal overgrowth, and temperature-induced bleaching (Williams et al. 2017), leading to their listing as a threatened species on the U.S. Endangered Species List in 2006 (National Marine Fisheries Service 2006). In 2015, the National Marine Fisheries Service published a formal Recovery Plan for *A. palmata* and *A. cervicornis*, which concluded that population recovery requires both conservation of genetic diversity and active restoration (National Marine Fisheries Service 2015). While coral restoration originally focused on fragmentation and asexual propagation of adult tissue (Young et al. 2012), the use of sexually-produced coral larvae for reef restoration is now widely practiced (Rinkevich 1995; Petersen et al. 2006; Omori et al. 2008; Rinkevich 2008; Omori 2011; Villanueva et al. 2012; Chamberland et al. 2015; Chamberland et al. 2016; dela Cruz and Harrison 2017; Pollock et al. 2017), and methods for larval propagation have been greatly improved (Vermeij et al. 2006; Vermeij et al. 2009; Marhaver et al. 2013; Marhaver et al. 2015; Chamberland et al. 2017; Pollock et al. 2017). Critically, larval propagation provides a mechanism to add tens of thousands of new genotypes *and* individuals to a coral population at one time while avoiding the logistical problems inherent in moving and fragmenting massive adult coral colonies.

According to microsatellite data (Baums et al. 2005a; Baums et al. 2005b), there are two genetically-isolated populations of *A. palmata* in the Caribbean: the western Caribbean population (including Panama, Mexico, Florida, the Bahamas, Navassa, and Mona Island) and the eastern Caribbean population (including the Virgin Islands, St. Vincent and the Grenadines, Bonaire, and Curaçao). A central, mixed genetic zone occurs between these two populations in Puerto Rico (Baums et al. 2005b). In Curaçao, we used cryopreserved sperm from Florida and Puerto Rico along with freshly-collected and cryopreserved sperm from Curaçao to conduct *in vitro* fertilization crosses, testing the reproductive compatibility across these genetic zones. Once larvae were produced, they were transferred to land-based aquaria in Florida for settlement and growth to examine the long-term survival of the AGF crosses (i.e., those that used sperm from Florida and Puerto Rico) in comparison to the local crosses (i.e., those that used sperm from Curaçao) created with both fresh and cryopreserved sperm. Larvae were moved to Florida prior to settlement so they could be acclimated to local conditions in the western Caribbean as early in the life cycle as possible. To date, no attempts have ever been made to use cryopreserved coral sperm to achieve assisted gene flow across genetically-isolated coral populations.

## Materials and Methods

### Study Sites and Gamete Collection

Two locations in Curaçao were chosen for spawning observations and gamete collection: Spanish Water (locally known as Spaanse Water; 12°4’13.11”N, 68°52’18.22”W) and the Curaçao Sea Aquarium (12°4’59.94”N, 68°53’42.47”W). Both reefs have large stands of *Acropora palmata* (Waitt Institute 2017). The reef at Sea Aquarium is known to have high overall genetic diversity (Baums et al. 2006) and Curaçao in general is known to have higher overall genetic diversity in *A. palmata* relative to other parts of the Caribbean (Baums et al. 2005a). In coordination with the full moons in late July and late August, divers surveyed between 25 and 100 *A. palmata* colonies per night. Observations were made from 2 days before the full moon to 11 days after the full moon in late July, and from 2 days before the full moon to 13 days after the full moon in late August, for a total of 30 nights of monitoring.

On each dive night, between 4 and 16 divers monitored colonies for at least 60 minutes, spanning the known spawning window of this species in Curaçao, beginning approximately 1 hour and 45 minutes after sunset. Divers examined colonies continuously for signs of setting (i.e., polyps holding egg-sperm gamete bundles in their mouths just prior to release). When setting was observed, the colonies were tented with weighted nylon mesh tents fixed to inverted plastic funnels in order to collect egg-sperm bundles in 50-mL conical centrifuge tubes (polypropylene, BD Falcon) affixed to the neck of each funnel, following methods previously developed by our team (Vermeij et al. 2006; Hagedorn et al. 2012a; Marhaver et al. 2013; Marhaver et al. 2015; Chamberland et al. 2016; Chamberland et al. 2017).

On shore, gamete collections from each colony were assessed for their suitability as either sperm donors or egg donors based on the volume of material produced; for each egg donor colony, we aimed to collect at least 2 mL of spawn so that replicate fertilization bins could be prepared containing at least 1,000 eggs per bin. The gamete samples that were chosen as egg donors were maintained in their original, closed collection tubes during transport back to the lab. Samples chosen for sperm pooling and cryopreservation were concentrated immediately upon arrival on shore by removing the majority of the seawater from the tubes using a new, sterile plastic transfer pipette for each tube, so that the remaining gamete bundles had approximately a 1:1 ratio of gamete volume to seawater volume in the tube. This ensured that after gamete bundles broke apart, the resultant sperm solution would be concentrated enough for successful cryopreservation. Highly-concentrated sperm samples can be difficult to collect, but high concentrations help to compensate for losses in viability due to freezing stress, and this allows for a smaller overall volume of the sperm solution and cryoprotectant to be used during *in vitro* fertilization. At this stage, our goal was to collect 5 mL of gamete bundles to 5 mL of seawater per sperm donor, but in some cases as little as 1 or 2 mL of spawn was used in order to achieve high sperm concentrations and to increase the overall number of donor genotypes that could be pooled. All gamete samples were transported about 40 minutes by car to the CARMABI Research Station for *in vitro* fertilization experiments.

In 2008 and 2016, sperm samples were collected and preserved from the western and central populations of *A. palmata*. For the central Caribbean samples, sperm was collected from Tres Palmas Marine Reserve in Rincón, Puerto Rico, in 2008. Sperm was pooled from five donor colonies and cryopreserved in sterile seawater (SSW; 0.2-µm impact filter, 47 mm, Millepore) with a final concentration of 5% dimethyl sulfoxide (DMSO, >99.5% purity, Sigma) as a cryoprotectant. For the western Caribbean samples, sperm was collected from Elbow Reef in Key Largo, FL, in 2016. For these samples, sperm were cryopreserved in SSW with a final concentration of 10% DMSO. In Florida, Elbow Reef is known to have low overall diversity and high clonality (Miller et al. 2016); therefore each sample was presumed to contain only one of two donor genotypes. All samples were held in storage under liquid nitrogen in the intervening years, then sent to Curaçao from the USDA National Animal Germplasm Program in Fort Collins, CO, via air using a liquid nitrogen dry shipper. Upon arrival in Curaçao, the temperature of the dry shipper was measured to be below −175°C, and it was immediately re-filled with liquid nitrogen. All samples were then held in storage under liquid nitrogen until immediately before they were thawed for *in vitro* fertilization in the laboratory.

### Egg Preparation and Screening

Egg-sperm bundles were allowed to break up with gentle or no agitation and eggs were then rinsed at least five times using filtered seawater (FSW; 47-mm-diameter GF/F filter, Whatman) in polycarbonate kitchen fat separators until the surrounding water was clear, indicating that residual sperm, plankton, and detritus had been removed. To avoid transferring any sperm from one parent colony to another, egg batches from each colony were kept in individual fat separators during the entire rinsing process, and separate seawater pouring beakers were used with each fat separator. Egg batches were then screened for the occurrence of self-fertilization. If eggs from a specific donor colony were observed undergoing primary cell cleavage, this indicated that fertilization had occurred within the collection tubes, and this material was not used for *in vitro* fertilization. Egg batches were observed for up to 4 hours after bundle breakup to avoid using eggs that were already fertilized. Trial experiments in Curaçao showed that the eggs remained viable for at least seven hours and *Acropora* eggs from the Pacific have been shown to remain viable for at least this long (Willis et al. 1997; Omori 2011).

### Sperm Preparation and Assessment

The concentrated samples chosen for sperm collection were gently agitated to break-up the gamete bundles, then the free sperm solution was removed from the bottom of the tube using a pipette and filtered through a clean cell strainer (70-µm nylon mesh, BD Falcon) to remove plankton, detritus, and any coral eggs carried over. Sperm samples were first kept separate by parent colony while they were assessed for motility under a phase microscope (Olympus BH2) and video system following method developed previously (Hagedorn et al. 2012a). To verify these data, additional motility and concentration data were collected by visual examination using a Leitz Orthoplan microscope with a phase contrast condenser and phase contrast objectives. Motility was scored by visual examination of 10-µL aliquots of the sperm solution (diluted 1:10 in clean FSW) that were spotted onto clean glass microscope slides and observed at 125×. Sperm concentrations were measured by direct observation using a cell counting chamber (sperm and bacteria counting chamber, Petroff-Hausser) at a total magnification of either 125× or 500×.

### Sperm Freezing and Thawing

Sperm samples were cryopreserved as described previously (Hagedorn et al. 2012a; Hagedorn et al. 2012b). Briefly, samples were kept as concentrated as possible, with the goal of achieving a sperm concentration above 1 × 10^9^ cells/mL. Samples in which total motility was at least 50% were pooled to create a mixed population of sperm from as many donor colonies as possible, with the goal of pooling material from at least 5 donor colonies per night to ensure pooled sperm contained substantial genetic diversity. The sperm concentration of each pooled sample was measured, then known volumes of the pooled sperm were diluted 1:1 (vol:vol) with freshly-prepared 20% DMSO in sterile seawater (SSW; 0.22-µm Sterviex HA syringe filter, Millepore). The 20% DMSO solution was added very gradually to the concentrated sperm with constant swirling to reduce exothermic heating, which can potentially damage the sperm. The sperm were allowed to equilibrate in the 10% DMSO for 10 min, then 1-mL aliquots were placed into cryovials (2.0 mL, externally threaded, Corning), which were capped, loaded into a custom-built cryofreezer, and frozen at 20 ± 2°C/minute (calculated as the slope of the line from −10 to -80°C) until they reached −80°C, at which time they were submerged directly in liquid nitrogen.

During nights when only a small number of colonies spawned in Curaçao, sperm was assessed, frozen, loaded into cryocanes, and maintained in liquid nitrogen for several days, then used on nights when sufficient eggs could be collected to perform the *in vitro* fertilization experiments. For the *in vitro* fertilization experiments conducted on 7 and 8 September 2018, the cryopreserved Curaçao sperm samples were frozen on 4 August 2018 (1 donor colony from Sea Aquarium) and 2 September 2018 (5 donor colonies from Spanish Water). These samples were pooled upon thawing, yielding a sperm pool for the CUR × CUR (frozen) crosses with material from six donor colonies. The inclusion of the sample from Sea Aquarium helped to ensure that the pooled sperm contained diverse genotypes. Tubes from these same batches of frozen Curaçao sperm were used for the *in vitro* fertilization trials on both 7 and 8 September 2018. The colonies at the Sea Aquarium site predominantly spawned on earlier nights in the lunar cycle, and with less synchrony overall. Further, weather conditions at this site made collecting difficult or impossible for some of the most prolific colonies. Therefore, these colonies were no longer monitored during the final four nights of the project. For the crosses described here, only fresh eggs from Spanish Water and frozen sperm from the Sea Aquarium and Spanish Water were used (Table 1).

**Table 1.** Summary of pooled sperm samples used to fertilize *A. palmata* eggs in Curacao. Sperm samples were collected and cryopreserved in three locations across the Caribbean. Sperm samples were pooled on the night of collection and/or after thawing to produce pools with as much genetic-diversity as possible. Freshly-collected sperm was also used for in vitro fertilization experiments. For the CUR (fresh) pool, material was mixed from either 4 or 5 males depending on the night of spawning and used on the same night it was collected. For the CUR (frozen) pool, material was mixed from two different sets of samples that were collected and frozen on different nights at different reefs (Sea Aquarium; 1 male, Spanish Water; 5 males). For the FL (frozen) pool, material was mixed from two different sample sets collected from different genotypes at Elbow Reef (“Green” and “Orange” genotypes). For the PR (frozen) pool, material was mixed from five males on a single night prior to freezing. Abbreviations: FL: Florida; PR: Puerto Rico; CUR: Curaçao.

| Sperm Pool ID for *in vitro* Fertilization | Sperm Collection Location and Reef(s) | Sperm Collection Date(s) | Number of Genotypes Pooled on Collection Date | Total Number of Genotypes Pooled on Fertilization Date | Date(s) Sperm Pool Used in Experiments |
| --- | --- | --- | --- | --- | --- |
| CUR (fresh) | CUR: Spanish Water | 7 Sept 2018 | 4 | 4 | 7 Sept 2018 |
| CUR (fresh) | CUR: Spanish Water | 8 Sept 2018 | 5 | 5 | 8 Sept 2018 |
| CUR (frozen) | CUR: Sea Aquarium | 4 Aug 2018 | 1 | 6 | 7 & 8 Sept 2018 |
|  | CUR: Spanish Water | 2 Sep 2018 | 5 |  |  |
| FL (frozen) | FL: Elbow Reef | 16 Aug 2016 | 1: "Orange Genotype" | 2 | 7 & 8 Sept 2018 |
|  | FL: Elbow Reef | 16 Aug 2016 | 1: "Green Genotype" |  |  |
| PR (frozen) | PR: Rincón | 7 Aug 2008 | 5 | 5 | 7 & 8 Sept 2018 |

To thaw cryopreserved sperm samples for *in vitro* fertilization, the cryovial was removed from liquid nitrogen and swirled gently in warm FSW (approximately 30°C) keeping the cryovial constantly moving (but without shaking) for about 2 minutes until the contents were completely thawed. The vial was then gently inverted to mix its contents, aliquots of sperm were quickly assessed for motility and concentration, and sperm was added to each fertilization container very gently using a micropipette to avoid placing heavy shear stress on the cells.

In Vitro *Fertilization Experiments*

On 7 and 8 September 2018 (nights 12 and 13 after the late August full moon), massive spawns were observed at Spanish Water, with approximately 75% of all colonies spawning during this window. On these nights, a series of large-scale *in vitro* treatments was performed using freshly-collected eggs from the Spanish Water colonies with four different pools of sperm. The four categories of crosses were: CUR × CUR (fresh sperm), CUR × CUR (frozen sperm), CUR × FL (frozen sperm), and CUR × PR (frozen sperm).

For all four categories of crosses, sperm was added to clear polystyrene containers (21.0 × 21.0 × 7.6 cm, ClearSeal clear hinged lid containers, Dart, Catalog #C90PST1) containing a starting water volume of 150 mL of FSW. Our goal was to aliquot 3,000 eggs per container and hold the number of eggs per container consistent between containers for each egg donor colony. In cases where donor colonies did not produce enough eggs to reach this target number, eggs were distributed evenly between containers for a total of 1,000 to 3,000 eggs per container.

*In vitro* crosses were designed to balance both experimental and conservation goals. First and foremost, we aimed to test whether AGF is possible in *A. palmata* to any degree, and if so, to produce as many AGF juveniles as possible with limited amounts of irreplaceable cryopreserved material. In a small-scale trial leading up to the mass spawning nights, we observed low overall fertilization in both the FL and PR sperm pools. Therefore, we added sperm to the large-scale *in vitro* crosses at two different concentrations; the low-sperm treatments were conducted to keep the final concentration of DMSO well below levels that can be toxic to sperm, while the high-sperm treatments boosted the overall sperm concentration to increase the number of encounters between sperm and egg, while potentially edging closer toward toxic levels of DMSO. For low-sperm treatments, we added 475 µL of the stock solution per container on both nights. For high-sperm treatments on 7 September, we added three times this amount (1425 µL per container). The only high-sperm treatment performed on 8 September was conducted in the CUR × PR (frozen) crosses using 950 µL of the stock solution.

Due to the natural variation in spawning volume on various nights when sperm samples were cryopreserved, there was also slight variation in the starting concentration of the cryopreserved sperm samples. For CUR (fresh sperm), starting sperm concentration was 5 × 10^8^ cells/mL and 6 × 10^8^ cells/mL, respectively, on 7 and 8 September 2018 (Table 2). For the frozen sperm samples, identical samples were used on both nights. Starting sperm concentrations were 6 × 10^8^ cells/mL for CUR (frozen sperm), 3.7 × 10^8^ for FL (frozen sperm), and 8 × 10^8^ for PR (frozen sperm).

**Table 2.** Summary of gamete handling steps and setup of *in vitro* fertilization crosses using sperm from genetically-isolated populations of *A. palmata*. Crosses were performed in Curaçao on 7 and 8 September 2018. Low: low-sperm treatments; 475 µL added on both nights. High: high-sperm treatments, 1425 µL added on 7 Sept, 950 µL on 8 Sept. For each pool of sperm, progressive motility was quantified upon thawing.

| Spawning Date | Sperm Pool ID | Number of Males | Progressive Motility (%) | Stock Sperm Conc. (cells/mL) | Final Sperm Concentration |  | Number of Eggs Used (Range) | Sperm Per Egg (Range) |
| --- | --- | --- | --- | --- | --- | --- | --- | --- |
|  |  |  |  |  | Low Sperm (cells/mL) | High Sperm (cells/mL) |  |  |
| 7 Sept 2018 | CUR Fresh | 4 | <25 to 50% | 5.0E+08 | 1.58E+06 | N/A | 1,000–2,000 | 118,750–237,500 |
| 7 Sept 2018 | CUR Frozen | 6 | 25% | 6.0E+08 | 1.90E+06 | 5.70E+06 | 1,000–2,000 | 142,500–855,000 |
| 7 Sept 2018 | FL Frozen | 2 | <25% | 3.7E+08 | 1.17E+06 | 3.52E+06 | 1,000–2,000 | 87,875–527,250 |
| 7 Sept 2018 | PR Frozen | 5 | <25% | 8.0E+08 | 2.53E+06 | 7.60E+06 | 1,000–2,000 | 190,000–1,140,000 |
| 8 Sept 2018 | CUR Fresh | 5 | 25 to 50% | 6.0E+08 | 1.90E+06 | N/A | 3,000 | 95,000 |
| 8 Sept 2018 | CUR Frozen | 6 | 25% | 6.0E+08 | 1.90E+06 | N/A | 3,000 | 95,000 |
| 8 Sept 2018 | FL Frozen | 2 | <25% | 3.7E+08 | 1.17E+06 | N/A | 3,000 | 58,583 |
| 8 Sept 2018 | PR Frozen | 5 | <25% | 8.0E+08 | 2.53E+06 | 5.07E+06 | 3,000 | 126,667–253,333 |

| Spawning Date | Female Colony ID | Spawn Volume | Eggs Allocated Per Bin | Water Per Bin | Number of Bins (Low Sperm + High Sperm) |  |  |  | Total Number of Bins | Total Number of Eggs Used | Colony Self-Fertilization |
| --- | --- | --- | --- | --- | --- | --- | --- | --- | --- | --- | --- |
|  |  |  |  |  | CUR Fresh | CUR Frozen | FL Frozen | PR Frozen |  |  |  |
| 7 Sept 2018 | F1 | 6 mL | 2,000 | 150 mL | 1 + 0 | 1 + 1 | 1 + 1 | 1 + 1 | 7 | 14,000 | NO |
| 7 Sept 2018 | F2 | 2.2 mL | 1,500 | 150 mL | 1 + 0 | 1 + 1 | 1 + 1 | 1 + 1 | 7 | 10,500 | NO |
| 7 Sept 2018 | F3 | 1.8 mL | 1,000 | 150 mL | 1 + 0 | 1 + 1 | 1 + 1 | 1 + 1 | 7 | 7,000 | YES |
| 7 Sept 2018 | F4 | 2.0 mL | 1,000 | 150 mL | 1 + 0 | 1 + 1 | 1 + 1 | 1 + 1 | 7 | 7,000 | YES |
| 8 Sept 2018 | F5 | 18 mL | 3,000 | 150 mL | 1 + 0 | 2 + 0 | 2 + 0 | 6 + 3 | 14 | 42,000 | NO |
| 8 Sept 2018 | F6 | 9 mL | 3,000 | 150 mL | 1 + 0 | 2 + 0 | 2 + 0 | 6 + 2 | 13 | 39,000 | NO |
| 8 Sept 2018 | F7 | 7 mL | 3,000 | 150 mL | 1 + 0 | 2 + 0 | 2 + 0 | 4 + 0 | 9 | 27,000 | YES |
| 8 Sept 2018 | F8 | 6 mL | 3,000 | 150 mL | 1 + 0 | 2 + 0 | 2 + 0 | 4 + 0 | 9 | 27,000 | NO |

Across both nights, final sperm concentration was between 1.17 × 10^6^ and 2.53 × 10^6^ cells/mL for the low-sperm treatments and between 3.52 × 10^6^ and 7.60 × 10^6^ cells/mL for the high-sperm treatments (Table 2). Overall, the final sperm-to-egg ratio in the *in vitro* fertilization experiments spanned the ratio considered optimal for corals (100,000:1) (Hagedorn et al. 2017). The final sperm:egg ratios in the experiments ranged from 87,000:1 to 1,140,000:1.

To continue monitoring egg batches for self-fertilization, replicated aliquots of eggs were taken from each donor colony and kept separate as no-sperm controls. Approximately 200 eggs were placed in 40 mL of FSW in 100-mm polystyrene Petri dishes, replicated three times per egg donor. These dishes were examined at multiple time points during the night to determine whether any cell division had taken place. These observations were not performed at the same large scale as the *in vitro* fertilization crosses because we prioritized the goal of producing as many juveniles of this threatened coral species as possible.

For each *in vitro* fertilization replicate, 1,000 to 3,000 thoroughly-rinsed eggs were placed in 150 mL of FSW in a polystyrene container, then either fresh or cryopreserved sperm was added by very slow pipetting and the solution was swirled gently every 1 to 2 minutes for approximately 10 minutes. Although fertilization may occur within minutes of adding sperm, cryopreserved sperm may have lower motility than fresh sperm, and the cryoprotectant in these samples can reduce sperm motility. Therefore, sperm-egg mixtures were left at this density for one hour with occasional swirling before any rinsing steps were started. After one hour, the volume in the containers was raised to 500 mL to dilute the cryoprotectant. During the next two hours, eggs and zygotes in every bin were rinsed several times with FSW to remove residual sperm, bacteria, and cryoprotectant, using a clean fat separator for each cross, then eggs and zygotes were transferred to clean containers containing new FSW as they began cell cleavage and embryogenesis.

Between 6 and 8 hours after sperm was added, fertilization success in each category of cross was assessed by sub-sampling between 40 and 150 embryos from each container. Fertilization was quantified during this window of time because the visual difference between unfertilized eggs and developing *A. palmata* embryos is most striking during gastrulation: unfertilized eggs remain round and intact while embryos undergoing gastrulation have the appearance of a “prawn chip” or “cornflake” (Okubo and Motokawa 2007). Furthermore, when coral eggs are fertilized with cryopreserved sperm, their time to first cleavage can be delayed by over an hour (Hagedorn and Carter 2016). Therefore, we assessed fertilization and development 6 to 8 hours after fertilization, rather than immediately after the onset of cleavage, as is done for other coral species (Levitan et al. 2004), to leave ample time for slower-developing embryos to proceed through cell division. The total number of unfertilized eggs and developing embryos were counted under a stereomicroscope (Nikon SMZ800) at between 10× and 63× magnification and the number of developing embryos was recorded. Fertilization percentages were then determined for each cross.

### Larval Propagation and Transport

After fertilization was quantified, all unfertilized eggs were removed from the containers by pipetting. The developing embryos were transferred into new FSW and new containers by pipetting, then embryos were distributed across additional containers as needed to maintain a density below 1 embryo per mL. This low density has been found to improve larval survival (Vermeij et al. 2006). Larvae were maintained in a dimly-lit, air-conditioned laboratory at 27°C with indirect natural light and a 12h:12h light:dark cycle with overhead fluorescent lights. Containers were held static with gentle agitation 4 to 5 times per day. Container changes and 95% water changes were performed every 24 to 48 hours by consolidating and rinsing larvae in a fat separator or by pipetting larvae into new containers. The number of swimming larvae in each treatment was assessed on 11 September 2018 (i.e., 3 and 4 days after spawning) in preparation for air transport to Miami, Florida, USA on 13 September 2018.

A variety of ultra-insulated coolers were tested using temperature loggers for their ability to maintain water temperatures between 27 and 28°C for at least 12 hours during air transport. Additionally, a variety of single-use and re-useable drinking water bottles were tested to identify which shape and material would cause the least shear stress and turbulence to the larvae. A simulated larval transport experiment was performed in which a small batch of *A. palmata* larvae was subjected to packaging, transportation, vibration, handling, and temperature stresses similar to those that would be experienced during air transit. Coolers produced by Ozark Trail (52 Quart High Performance Cooler; interior dimensions 58 × 29 × 28 cm L × W × H) and Pelican (Pelican Elite 30; interior dimensions 37 × 26 × 28 cm L × W × H), and 1.5 L HDPE clear plastic bottles with narrow necks (Lovers Ice Water, Curaçao; 9 × 28 cm W × H) were chosen to transport the larvae. Bottles were kept sealed prior to shipping the larvae, and only opened immediately before use. Fresh water was decanted and bottles were rinsed once with FSW.

Larvae were consolidated in freshly-prepared FSW using clean glass and plastic transfer pipettes. Approximately 150 to 1000 larvae were packaged per bottle, depending on the relative rarity of each larval cohort (i.e., corals from AGF crosses were packed at lower density to maximize survival). For all bottles, larval density was kept below 1 larva per mL FSW to maintain high dissolved oxygen concentrations and discourage bacterial growth. Bottles were filled so that less than 1 cm of vertical air space remained in the neck, which reduced water sloshing and subsequent shear stress on the larvae. Lids were tightened only in the final 10 minutes before closing the coolers. Each cooler contained 12 bottles. Excess space in the Ozark Trail cooler was filled with foil-lined, ultra-insulating bubble wrap to stabilize the larvae and further buffer against temperature changes.

Larvae were packaged into the bottles beginning at 03:00 Eastern Time (ET) on the day of transport. Air travel began at 07:00, arrival in Miami was completed by 11:00, customs clearance was completed by 12:00, and ground transportation was completed to the destination facilities by 15:00 ET. Thus, larvae spent approximately 12 hours in the bottles. Upon arrival, larvae could be seen actively swimming in the bottles. A total of 60 L of water and over 20,000 larvae were transported using four coolers. At the two settlement facilities, Mote Marine Lab and The Florida Aquarium Center for Conservation, larvae were carefully poured into holding containers (1-L polystyrene clamshell containers) to recover from shipment. The number of larvae from each bottle was estimated and compared to the number of larvae shipped.

### Larval Settlement and Grow-out at Mote Marine Lab

Over 20,000 coral larvae were transported by air to Florida, where the larval cohort was divided for settlement and grow-out at Mote Marine Laboratory and The Florida Aquarium. At Mote, larval settlement was carried out in static 19-L glass aquaria containing seawater from a well system. Seawater exiting this well (pH 7.5) was immediately aerated to off-gas carbon dioxide and hydrogen sulfide, filtered through a moving bed biological filter containing SWX media (Sweetwater) followed by a sand filter (Pentair Aquatic Ecosystems), then passed through a pleated filter (100 µm, Pentair Aquatic Ecosystems) to remove sediment before being introduced to settlement aquaria (pH 8.0, Salinity 37 ppt). Aquaria were maintained in an indoor, temperature-controlled wet lab at approximately 27°C. Aquaria were also halfway submerged in flow-through fiberglass raceways measuring 2.5 × 1.0 × 0.3 m to act as a second temperature control failsafe. Swimming larvae were added to the aquaria each containing 12-20 replicate ceramic substrates (Ceramic Coral Frag Plugs, Boston Aqua Farms) measuring 3 cm in diameter.

To estimate larval numbers, larvae were transferred to a vessel with a known volume and the total volume of seawater was raised to 2 L. The water was gently mixed to disperse the larvae uniformly in the container, then a glass pipette (10 mL, Pyrex) was used to draw up a volume of seawater and larvae. The pipette was viewed under a stereomicroscope (American Optical) at 10× magnification and larvae were quantified by counting the number present in 3 separate, 1-mL sections of the pipette. This entire process was then repeated two more times and the resulting values were averaged to arrive at an estimate of larval number per mL. The larvae were then aliquoted in know numbers to settlement tanks.

Settlement was fostered by sprinkling ground pieces of live crustose coralline algae (CCA) onto the tops of each substrate. The CCA used was a mixture of unidentified species cultured in Mote’s land-based coral farm in the same location. Importantly, the addition of this mixture of diverse CCA encouraged most of the swimming larvae to settle onto the tops of each substrate rather than onto the undersides of the substrates or onto glass surfaces in the tanks. Given the high conservation value of the two larval cohorts produced from AGF sperm (i.e., larvae from the CUR × FL and CUR × PR crosses), and to maximize the number of settlers obtained from these two groups, extra effort was taken to foster their settlement by keeping them at a lower density compared to the other larval cohorts. For these AGF cohorts, approximately 250 larvae were added per settlement tank. Similarly, larvae from the CUR × CUR (frozen) cross were given higher priority than larvae from the CUR × CUR (fresh) cross because larvae were more abundant in the latter cross, and the former has conservation value as the largest coral cohort ever produced through cryopreservation. Thus, approximately 750 larvae were allocated per settlement tank for the CUR × CUR (frozen) cross and approximately 1700 larvae were allocated per settlement tank for the CUR × CUR (fresh) cross. After larvae had settled, the substrates were moved to a dimly-lit, indoor, flow-through raceway (same dimensions as above) for post-settlement care and grow-out. The raceway was fed by flow-through seawater at 1.0 L/minute and 6-7.5-cm air stones were used to provide water circulation within the raceway.

At approximately 1.5 months post-settlement, all settled corals were moved to a clean, outdoor grow-out raceway with the same dimensions and number of air stones as above, fed by unfiltered seawater at a rate of 4.0 L/minute at 23-27°C and pH 8.0. The settlement substrates were marked according to each cross on the bottom of each pedestal with permanent marker covered by extra thick cyanoacrylate super glue gel (Bulk Reef Supply) and placed on separate plastic egg crate racks. Algal fouling was reduced by elevating the racks 3 cm above the raceway floor with PVC legs and with the addition of the intertidal snail *Batillaria minima* (at a density of approximately 700 snails/2.5-m raceway). Each day, the bottom of the raceway was siphoned to remove detritus and grazers were collected and redistributed evenly across the raceway. Substrates were monitored daily and undesirable fouling organisms that recruited to each substrate were smothered with super glue before they were able to grow and proliferate to other substrates. These methods helped to encourage growth by the CCA on pedestals. Although CCA can at times overgrow coral settlers, this group of encrusting organisms is easier to monitor and manage compared to other coral competitors such as turf algae, fleshy algae, and cyanobacteria. As needed, egg crate racks, air stones, and the raceway were removed and replaced with clean backups whenever surfaces became heavily fouled. Finally, for all corals in the AGF crosses, substrates were monitored for potential overgrowth by fast-growing CCA. Any rapidly-encroaching CCA was removed from around each recruit using a scalpel (X-Acto knife, Elmer’s Inc.) to prevent potential smothering of the coral polyps.

### Larval Settlement and Grow-out at The Florida Aquarium Center for Conservation

Upon arrival at The Florida Aquarium, larvae were poured gently into polystyrene clamshell containers and water from the settlement aquarium was mixed to a final ratio of 50% shipping water to 50% settlement aquarium water. Temperature and salinity were matched within 0.5°C and 1 ppt between the shipping water and the settlement aquarium. Larvae were then transferred by gentle pouring into the settlement containers. Settlement containers were constructed out of 17-L polyethylene dish pans with four 7.6 cm holes drilled into the sides. Holes were covered with 150-µm nylon mesh attached to the bin with food-grade silicone sealant. Settlement containers were placed on a PVC rack inside of a well-established, 2270-L recirculating aquarium system with live rock, a deep sand bed, and protein skimming. A pump and valve manifold supplied a slow trickle of water through each settlement bin from the main aquarium.

Larvae from the AGF crosses (i.e., CUR × FL and CUR × PR) were placed at a lower density in the settlement bins, ranging from approximately 250 to 550 per bin. Larvae from the Curaçao fresh and frozen sperm crosses (i.e., CUR × CUR (fresh) and CUR × CUR (frozen)) were placed at a higher density of larvae per bin, at 1000 and 1500 larvae per bin, respectively. Settlement tiles were smooth ceramic squares (Ceramic Reef Squares, Boston Aqua Farms) that were conditioned for three months in recirculating coral aquaria with small colonies of *Acropora cervicornis* and *A. palmata.* In addition, fragments of CCA (tentatively identified as *Hydrolithon boergesenii*) were ground using a mortar and pestle and sprinkled into the settlement aquaria. In order to provide a source of free-living coral symbionts and to provide additional settlement cues for the larvae, a small amount of sediment from well-established aquaria was also added to the settlement aquaria.

Within one week of settlement, settled polyps were counted and transferred into two separate recirculating aquarium systems located in a climate-controlled greenhouse. One system consisted of a single shallow raceway (2.4 m × 0.9 m × 0.3 m) and a sump (1700 L total) and the second system consisted of two shallow raceways (1.5 m × 0.6 m × 0.4 m) and a sump (1325 L total). Each system contained live rock, a deep sand bed (10-12 cm deep), and several fragments of *A. cervicornis* to serve as sources of additional symbionts. Each system contained a protein skimmer, titanium immersion heaters, titanium chilling coils, and a media reactor with activated carbon. Temperature was set at 25 ± 0.5 °C. Aquaria were covered with shade cloth so that they received 50-75 µmol m^-2^ s^-1^ of photosynthetically active radiation (PAR) at solar noon. Algae that grew around coral settlers was manually removed as needed using aluminum clay needle tools and curved, stainless steel tweezers. To reduce algal fouling, both aquaria were stocked with small herbivorous snails (*Batillaria minima, Cerithium lutosum*, and juvenile *Lithopoma americanum*).

At approximately two weeks post-settlement, a visible tissue loss syndrome was observed beginning in the CUR × CUR (frozen) cohort, which rapidly spread to other groups of settlers. To halt further losses, all settlers at the Florida Aquarium Center for Conservation were given a 10-day bath in ampicillin dosed at 100 µg mL^-1^ based on previous studies (Vermeij et al. 2009; Sweet et al. 2014). Recruits were moved into a single system and isolated from filtration, and a 100% water change was conducted daily with a re-dose of antibiotics each day. This treatment stopped the spread of the tissue loss syndrome, after which recruits were moved into a new aquarium system with cycled live rock and returned to circulation with filtration. After one month, PAR levels were increased to 150 µmol m^-2^ s^-1^.

## Results

### Spawning Timing and Behavior

In total, divers surveyed *A. palmata* colonies at two separate sites during 14 nights in late July and early August, and 16 nights in late August and early September. Although spawning in this species can generally be narrowed down to a period of less than a week, its spawning times in Curaçao tend to be less predictable than in other locations in the Caribbean (M. Vermeij, V. Chamberland, personal observation), and the timing of the full moons relative to the solar calendar in 2018 made spawning less straightforward to predict. In total, we observed at least one *A. palmata* colony spawn on night 8 after the 27 July full moon and on the following nights surrounding the 26 August full moon: −2, −1, +4, +5, and +7 through +13. The largest mass spawns (during which more than 50% of colonies spawned) occurred on 7 and 8 September 2018, which were nights +12 and +13 after the late August full moon.

### Sperm Motility

Sperm motility was quantified for each sample collected in Curaçao, and for each thawed sample from Curaçao, Puerto Rico, and Florida that was used in this study. In fresh sperm samples collected in Curaçao, total mean motility ranged from 25 to 50% and progressive motility ranged from 26 to 38% (Table 2). Sperm in these samples remained motile for at least 6 hours. These sperm motility values were within the range observed in *A. palmata* from other locations, including the Florida and Puerto Rico samples collected in 2016 and 2008, respectively (Hagedorn et al. 2012a). The post-thaw motility for the frozen samples from Curaçao was approximately 25%. Post-thaw motility for the frozen samples from Florida and Puerto Rico was between 0 and 25% (Table 2.).

### Fertilization and Development of Embryos Produced with AGF Sperm

Four categories of *in vitro* fertilization crosses were performed using freshly-collected *A. palmata* eggs from Curaçao (Figure 1). The crosses were designated: CUR × CUR (fresh sperm), CUR × CUR (frozen sperm), CUR × FL (frozen sperm), and CUR × PR (frozen sperm). On 7 and 8 September 2018, we mixed *A. palmata* sperm from the four different sperm pools with eggs from 8 different colonies (Table 2). Despite screening and removing material that underwent self-fertilization prior to the addition of sperm, eggs from 3 of the 8 colonies *also* underwent self-fertilization after crosses were mixed. This was identified by observing cell division rates of 50-95% in no-sperm controls, compared to the cell division rates normally observed in no-sperm controls (0-1%). The fertilization data from these three colonies was removed from all calculations of mean fertilization rate by sperm pool, leaving data from five colonies that did not self-fertilize. The larvae produced from these three uncontrolled crosses were kept alive because there was a chance that a small fraction of the juveniles produced in these containers would still be the result of fertilization by the AGF sperm that was added. This can only be determined at a later date through genetic analysis.

**Figure 1.**
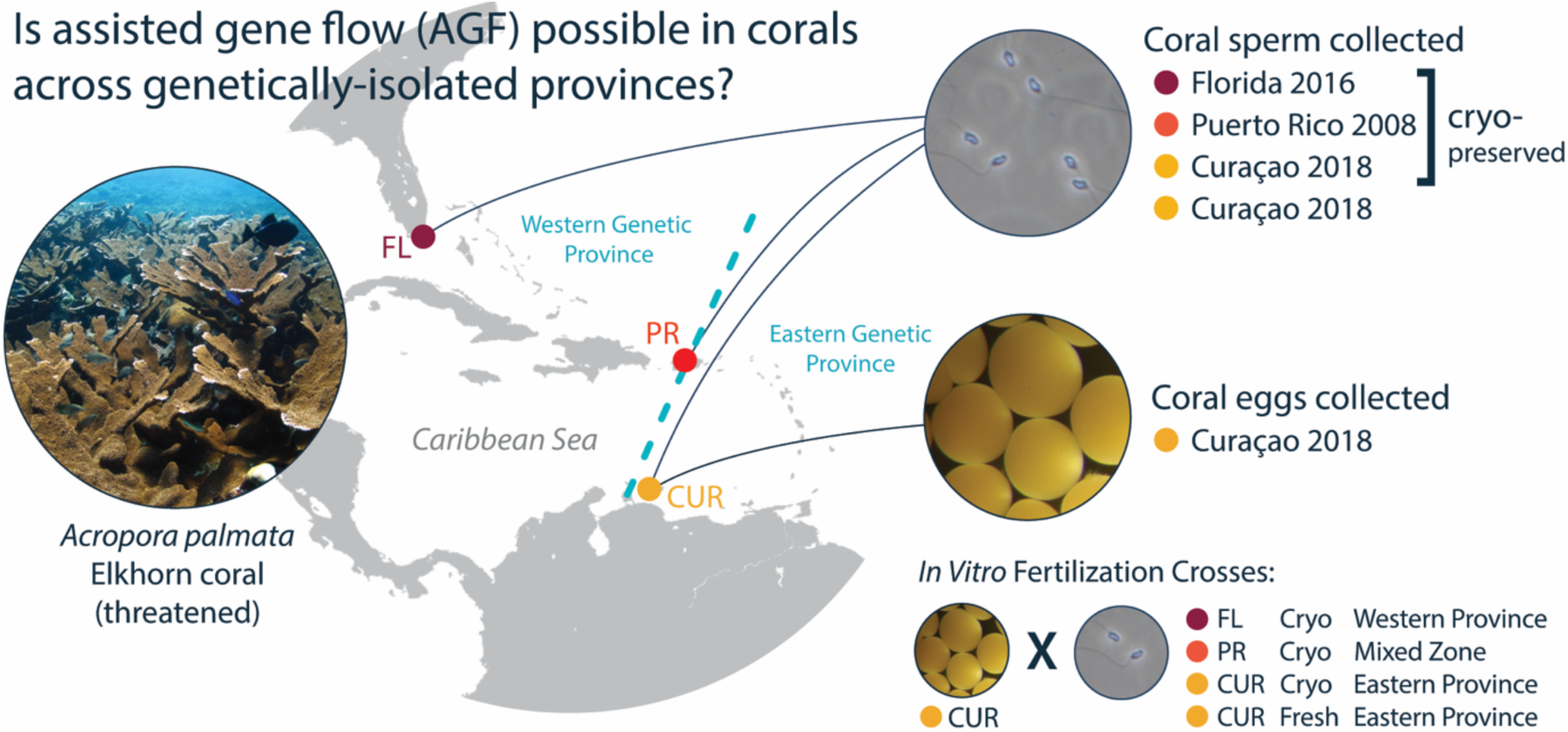
Summary of sperm collections and *in vitro* crosses conducted using cryopreserved material from genetically-isolated populations of *A. palmata* in the Caribbean to test the feasibility of assisted gene flow (AGF). Sperm samples from Curaçao, Florida, and Puerto Rico were cryopreserved and moved to Curaçao, then used for *in vitro* fertilization experiments with freshly-collected eggs from Curaçao. Freshly-collected sperm from Curaçao was used for comparison. These crosses represent the first attempts to fertilize *A. palmata* using cryopreserved sperm and the first trials of assisted gene flow (AGF) across genetically-distinct populations of Caribbean corals.

For the remaining crosses (in which no self-fertilization occurred: Table 3; Figure 2), mean fertilization rate was 91 to 99% for CUR × CUR (fresh sperm) crosses, 37 to 82% for CUR × CUR (frozen sperm) crosses, 3 to 19% for CUR × FL (frozen sperm) crosses and 0 to 24% for CUR × PR (frozen sperm) crosses. The CUR × CUR (fresh) crosses achieved fertilization rates of 91 to 99%. The CUR × CUR (frozen) crosses from 7 September achieved fertilization rates of 80 to 82%. The same category of crosses on the following night achieved 37 to 60% fertilization. On both nights, the AGF crosses had lower rates of fertilization: CUR × FL (frozen) crosses had fertilization rates of 10 to 19% and 3 to 18% on 7 and 8 September, respectively, while CUR × PR (frozen) crosses had 2 to 24% and 0 to 2% fertilization on those same nights.

**Table 3.** Fertilization rates for *A. palmata* eggs crossed with sperm from genetically-isolated populations. Crosses were performed in Curaçao on 7 and 8 September 2018. Data represent the percentage of eggs/zygotes that were actively developing (i.e., undergoing embryogenesis) approximately 6 hours after sperm addition. No Sperm: Eggs were kept in FSW after cleaning and no sperm was added from any source. Low: low-sperm treatments; 475 µL added on both nights. High: high-sperm treatments, 1425 µL added on 7 September, 950 µL on 8 September. N/D: No Data: Negative controls for Female F5 were not replicated due to a lab pipetting oversight in the early morning hours. A fertilization score was made from the egg stock at 7 AM instead. N/A: Not Applicable: High-sperm treatments were not performed for all sperm pools on all nights. Data are show here by sperm treatment. In contrast, data are shown in Figure 2 as averages across both low and high sperm treatments for each egg donor colony.

| Female Colony ID | Female Spawning Date | Sperm Pool |  |  |  |  |  |  |  |  |  |
| --- | --- | --- | --- | --- | --- | --- | --- | --- | --- | --- | --- |
|  |  | No Sperm 1 | No Sperm 2 | No Sperm 3 | CUR Fresh Low | CUR Frozen Low | CUR Frozen High | FL Frozen Low | FL Frozen High | PR Frozen Low | PR Frozen High |
| F1 | 7 Sept 2018 | 0.0% | 0.7% | 0.5% | 99% | 82% | 82% | 10% | 18% | 2% | 2% |
| F2 | 7 Sept 2018 | 0% | 0% | 0% | 94% | 82% | 80% | 18% | 19% | 24% | 8% |
| F5 | 8 Sept 2018 | 0.1% | N/D | N/D | 91% | 58% | N/A | 18% | N/A | 1% | 2% |
| F6 | 8 Sept 2018 | 0% | 0% | 0% | 93% | 37% | N/A | 3% | N/A | 0% | 0% |
| F8 | 8 Sept 2018 | 0% | 0% | 0% | 95% | 60% | N/A | 10% | N/A | 0% | 0% |

**Figure 2.**
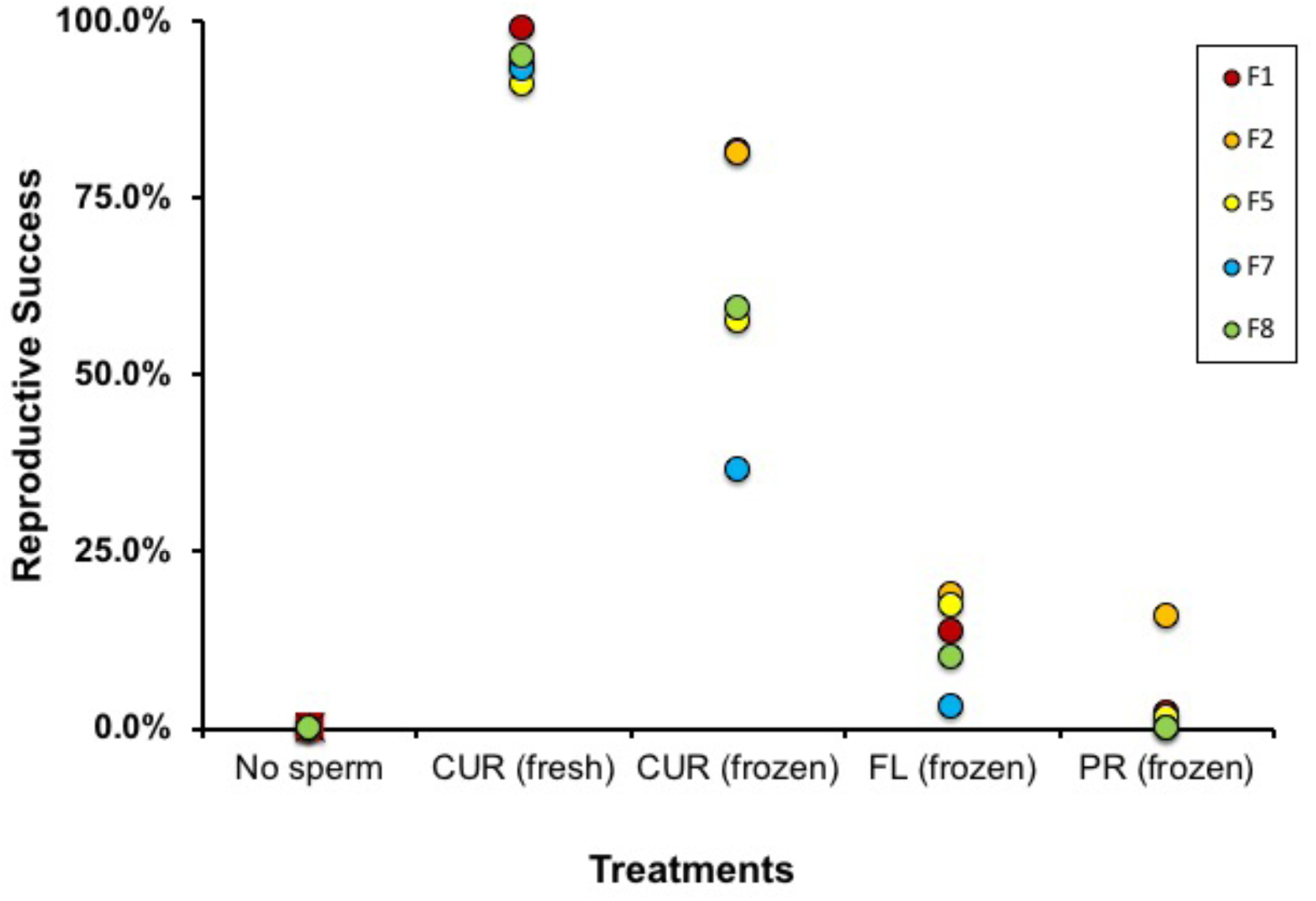
Mean fertilization percentage for *A. palmata* eggs fertilized by one of four sperm pools collected from genetically-isolated populations. All crosses were performed with eggs collected in Curaçao (CUR) and with fresh or cryopreserved (frozen) sperm from Curaçao (CUR), or cryopreserved sperm from Florida (FL) or Puerto Rico (PR). Colonies spawned in Curaçao on 7 and 8 September 2018. Egg donors are numbered separately by donor colony: F1, F2, F5, F7, F8 (See Table 2). Because no consistent effect of sperm concentration was observed, data for each egg donor were averaged across sperm density treatments prior to plotting.

During embryogenesis, the larvae produced with AGF sperm developed normally. No abnormal embryos were observed except in a small number of containers that were rinsed too aggressively during a fragile stage of embryogenesis. Overall, we observed no evidence that eggs fertilized by cryopreserved sperm were compromised in any way during early cell division and embryogenesis.

### Juvenile Settlement, Survival, and Symbiont Acquisition

No noticeable loss in larvae was observed during transport and no dead larvae were found during unpacking, suggesting that all of the transported larvae survived in good condition. After approximately 1 week in settlement containers, 40 and 45% of the larvae had successfully completed settlement and metamorphosis at Mote Marine Lab and The Florida Aquarium, respectively, yielding over 11800 settled primary polyps in total. The overall settlement rate in the AGF larvae was 54%. The overall settlement in the CUR × CUR (fresh) and CUR × CUR (frozen) crosses was 46% and 45%, respectively (Table 4).

**Table 4.** Summary of larval settlement and survival rates for *A. palmata* juveniles produced through *in vitro* fertilization with sperm from genetically-isolated populations. Larvae were reared in Curaçao and shipped to two facilities in Florida for settlement and long-term grow-out. Approximately equal numbers of larvae were shipped to each facility, which employed custom, in-house methods to foster larval settlement and post-settlement growth. Larval cohorts were handled using similar methods, but increased care was directed toward AGF larvae, i.e., larvae that resulted from crosses between genetically-distinct populations of the Caribbean (CUR × FL and CUR × PR cohorts). The large decline in survivorship from settlement to month 1 was primarily caused by an outbreak of bacterial disease.

| Category of gamete cross:<br>EGG × SPERM (Sperm Type) | Number of<br>Larvae | Initial Number<br>of Settlers | Settlement<br>Rate | Number of<br>Settlers at 1<br>month | Survival<br>Rate at 1<br>Month |
| --- | --- | --- | --- | --- | --- |
| The Florida Aquarium |  |  |  |  |  |
| CUR × CUR (Fresh Sperm) | 3450 | 1847 | 54% | 964 | 52% |
| CUR × CUR (Frozen Sperm) | 4000 | 2111 | 53% | 292 | 14% |
| CUR × FL (Frozen Sperm; AGF) | 1107 | 663 | 60% | 367 | 55% |
| CUR × PR (Frozen Sperm; AGF) | 270 | 100 | 37% | 63 | 63% |
| CUR × Unknown (Self-Fertilization) | 6700 | 2257 | 34% | 828 | 38% |
| Location Totals | 15527 | 6978 | 45% | 2514 | 36% |
| Mote Marine Lab |  |  |  |  |  |
| CUR × CUR (Fresh Sperm) | 3400 | 1258 | 37% | 1205 | 96% |
| CUR × CUR (Frozen Sperm) | 4000 | 1466 | 37% | 1293 | 88% |
| CUR × FL (Frozen Sperm; AGF) | 1100 | 584 | 53% | 636 | 109% |
| CUR × PR (Frozen Sperm; AGF) | 270 | 133 | 49% | 127 | 95% |
| CUR × Unknown (Self-Fertilization) | 3500 | 1431 | 41% | 667 | 47% |
| Location Totals | 12270 | 4872 | 40% | 3928 | 81% |
| Both Institutions |  |  |  |  |  |
| CUR × CUR (Fresh Sperm) | 6850 | 3105 | 46% | 2169 | 70% |
| CUR × CUR (Frozen Sperm) | 8000 | 3577 | 45% | 1585 | 44% |
| CUR × FL (Frozen Sperm; AGF) | 2207 | 1247 | 57% | 1003 | 80% |
| CUR × PR (Frozen Sperm; AGF) | 540 | 233 | 42% | 190 | 82% |
| CUR × Unknown (Self-Fertilization) | 10200 | 3688 | 52% | 1495 | 41% |
| Grand Total: Frozen Sperm | 10747 | 5057 | 47% | 2778 | 55% |
| Grand Total: AGF Sperm | 2747 | 1480 | 54% | 1193 | 81% |
| Grand Total: All Crosses | 27797 | 11850 | 43% | 6442 | 54% |

Settlers were first observed taking up symbiotic dinoflagellates (*Symbiodiniaceae* spp.), either from the water column or from the ground CCA, within 5-10 days post settlement (Figure 3; Panels A and B). Within 1 month, approximately 75% of settlers were well-infected with symbionts, evidenced by the dark orange-brown color of their tissues (Figure 3; Panels C-F).

**Figure 3.**
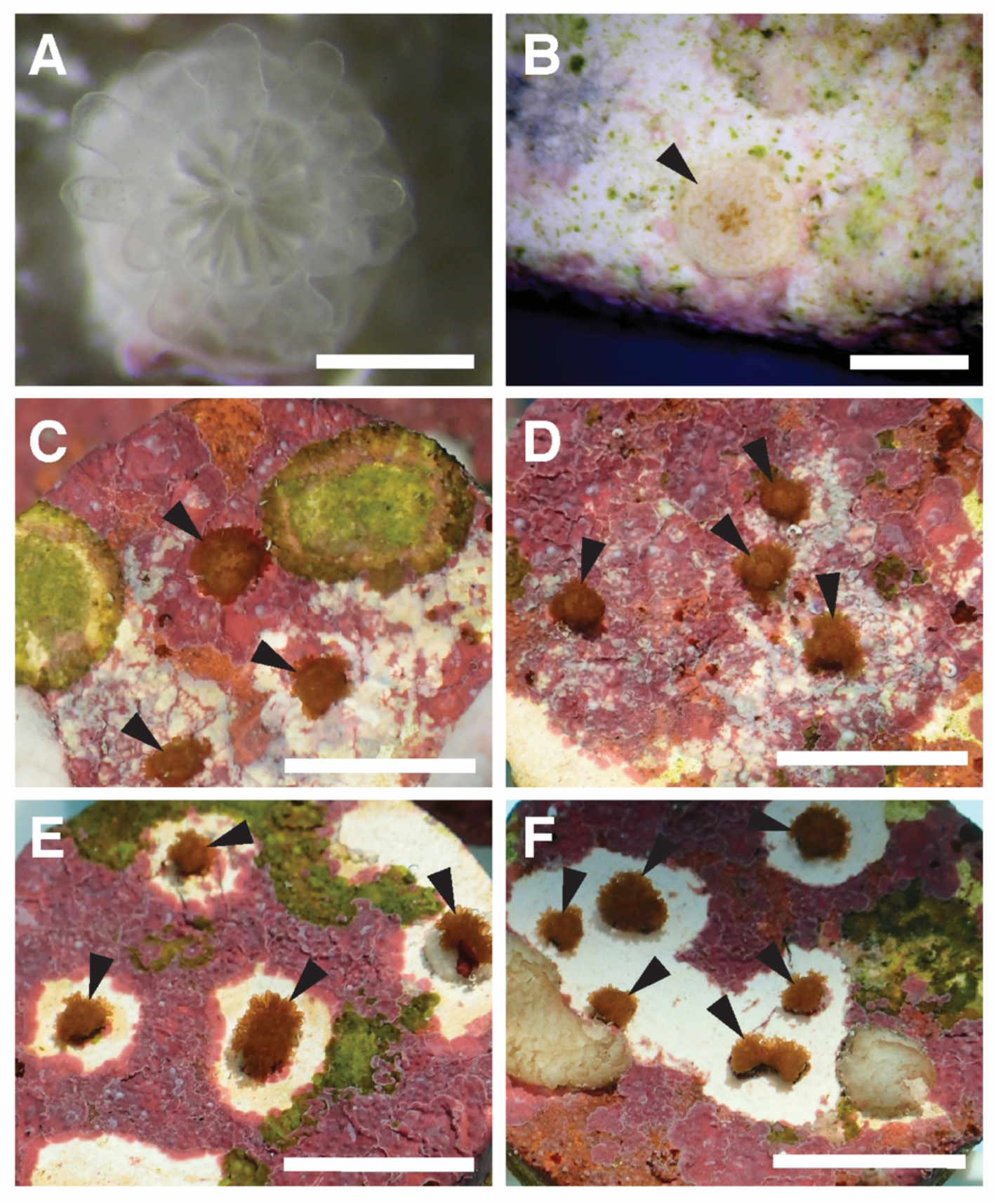
Successful settlement, symbiont acquisition, and growth of juvenile *A. palmata* corals produced from cryopreserved sperm from genetically-isolated populations. All crosses were performed with eggs collected in Curaçao (CUR) and with either fresh or cryopreserved (frozen) sperm from Curaçao (CUR), Florida (FL), or Puerto Rico (PR). A) Three-week-old, settled juvenile from the CUR × CUR (frozen) cross, showing successful metamorphosis and skeletogenesis. Early stages of symbiont acquisition can be seen as faint, darker spots in the tentacles. Scale bar = 0.25 cm. B) Three-week-old, settled juvenile from CUR × FL (frozen) crosses (black arrowhead), showing increased symbiont density (brown tentacles) and skeletal growth. Scale bar = 0.5 cm. C?F) Six-week-old, settled juveniles (black arrowheads) from all four categories of crosses, showing high density of symbionts (dark red-brown color), budding of new polyps, and skeletal growth. C) CUR × CUR (fresh) cross. D) CUR × CUR (frozen) cross. E) CUR × FL (frozen) cross. F) CUR × PR (frozen) cross. For panels C?F, Scale bars = 1 cm. White areas surrounding the settlers represent zones where the coral juveniles are inhibiting the growth of CCA or where CCA was removed with a scalpel to prevent overgrowth. No differences are apparent between the settlers from the four crosses in gross morphology, development, or relative health.

At one month, the numbers of surviving settlers were counted in each of the four crosses (Table 4). As a total of all four categories, overall settler survival was 81% at Mote Marine Lab. Survival at The Florida Aquarium was lower due to the outbreak of a tissue loss syndrome, which was arrested by antibiotic treatment but nevertheless caused heavy losses in the CUR × CUR (frozen) settlers and approximately 30% of the settlers in all other groups. Overall settler survival at The Florida Aquarium was 36%. However, the tissue loss syndrome occurred primarily in the larger population of settlers from the CUR × CUR (frozen) crosses, which had a one-month survival rate of 14%. The remaining cohorts at Florida Aquarium had one-month survival rates of 37 to 63%. Across both institutions, the one-month survival rate of all juveniles was 54%, the one-month survival rate of juveniles produced from cryopreserved sperm was 55%, and the survival rate of juveniles from the Florida and Puerto Rico AGF crosses was 81% (Table 4).

## Discussion

### First Demonstration of AGF in Caribbean Corals

Given the conservation importance of *A. palmata*, and the conservation promise of assisted gene flow, we sought to test whether cryopreserved *A. palmata* sperm from populations in the western and central Caribbean could be used to successfully fertilize *A. palmata* eggs from the eastern Caribbean. In doing so, we tested for the first time whether genetically-isolated populations of this threatened species are in fact reproductively compatible. If so, these genetically isolated populations would have the potential to be cross-bred for conservation purposes. Furthermore, we tested for the first time whether cryopreserved sperm from *A. palmata* can successfully fertilize *A. palmata* eggs at all, as there have been no previous attempts to perform *in vitro* fertilization using cryopreserved material in this threatened coral species. Finally, by conducting *in vitro* fertilization in corals using cryopreserved material, we aimed to demonstrate the potential for cryopreservation to enhance endangered species conservation at increased scales by producing the largest living wildlife population ever created from cryopreserved material. Although AGF experiments have been successfully conducted on the Great Barrier Reef with live material by moving whole coral colonies (Dixon et al. 2015), the research we present here is the first example of using cryopreserved material to cross-breed two genetically-distinct coral populations, and the first test of AGF using any Caribbean coral species. This proof-of concept experiment opens a wide range of options for future studies and conservation initiatives using cryopreservation of coral sperm.

### Utility of Cryopreserved Material in Coral Conservation

When cells are frozen and banked properly, they can retain viability for years, as shown by the ten-year-old sperm samples from Puerto Rico that were used to achieve fertilization in this study. Thus, cryopreservation is a valuable means to safeguard genetic diversity of existing species and populations while it still exists. Genome resource banks containing cryopreserved material can help to 1) preserve and protect large gene pools that can be used to rescue genetically-depleted populations, 2) transport genetic materials among widely-dispersed living populations relatively easily and inexpensively, 3) prolong the effective generation times of organisms with short generation times, and 4) improve the speed and effectiveness of research on germplasm and animal reproduction. However, cryopreservation does stress the cells involved. In corals, this can result in lower sperm motility of thawed samples compared to fresh samples, a longer time until cell cleavage begins, and potentially higher variability in reproductive success from trial-to-trial (Hagedorn et al. 2012a; Hagedorn et al. 2017). These effects of cryopreservation appear to be restricted to the early stages of fertilization and the initiation of embryogenesis. Thus far, we have not observed any differences in the settlement and post-settlement growth of juvenile corals produced with cryopreserved sperm as compared to those produced with fresh sperm (Hagedorn et al. 2017).

### Challenges of Controlling Coral Fertilization

This proof-of-concept project revealed a range of logistical and biological issues that must be considered in order to perform successful cryopreservation and controlled *in vitro* fertilization with spawning corals. First, by observing *A. palmata* colonies for 30 total nights across two separate lunar cycles, we found that spawning by this coral species is not always synchronized across colonies in one or two “mass” spawning nights. Instead, there were multiple periods during which small levels of spawning were observed prior to the mass spawning that finally occurred. This variability has potentially negative consequences for natural fertilization rates in any coral population that is prone to strong Allee effects. Furthermore, these observations are important for future restoration programs to consider when planning the duration and level of effort for spawning operations.

A second challenge encountered in this study was the propensity for eggs collected from individual *A. palmata* colonies to undergo fertilization before any sperm samples could be added. This posed a major challenge to controlling crosses all the way through the process of *in vitro* fertilization. Indeed, *A. palmata* and the vast majority of hermaphroditic spawning corals are generally understood and observed to be self-incompatible at the genotype level. However, *A. palmata* is a very long-lived species, with some genotypes estimated to be hundreds of years old (Irwin et al. 2017) and it is highly prone to fragmentation. Moreover, *A. palmata* larvae have been observed to settle gregariously (i.e., in closely-packed groups). The fact that we observed self-fertilization at the colony level suggests that one or two different mechanisms may be at work. First, individual colonies may be chimeras of more than one genotype, either as a result of gregarious larval settlement or the fusion of fragments from different genotypes (Schweinsberg et al. 2015). If these two (or more) genotypes are reproductively compatible, we would expect their gametes to fertilize one another within the gamete collection tube. Second, somatic mutations accumulate in adult corals, especially in relatively old genotypes (Irwin et al. 2017); such mutations could cause the loss of functional self-incompatibility genes. This would lead to self-fertilization by gametes from only this single genotype in the collection tube. These somatic mutations might also be heritable, producing juveniles that themselves are self-compatible. We took a number of measures to avoid and eliminate self-fertilizing material in this study. For example, because some colonies spawned across multiple consecutive nights during this study, we were able to identify colonies and stands of *A. palmata* that underwent self-fertilization to avoid collecting from them on subsequent nights. Further, we observed eggs for as long as possible prior to sperm addition to avoid using self-fertilized material. Nevertheless, a subset of eggs used in our crosses underwent self-fertilization. This represented a barrier to producing a greater number of AGF juveniles in the current study.

### Utility of coral assisted gene flow relative to other interventions

As a form of both managed breeding and assisted migration, AGF has been proposed as a reasonable and realistic intervention for coral populations, especially given the rapid pace of global change and the slow turnover of coral populations (National Academies of Sciences Engineering and Medicine 2018). Although in general, gene flow into a population can introduce genes unsuitable for a local environment and thus reduce a population’s degree of local adaptation (a phenomenon known as outbreeding depression), the risk of this is considered low in corals given their extant genetic diversity and population genetic structure. Importantly, AGF does not involve any removal of genes or introduction of genes, nor does it require knowledge of the genetics underlying environmental adaptation, all of which limit the practicality and applicability of more extreme coral interventions such as gene editing. In addition, AGF can be conducted without selecting for individual phenotypes at the expense of others, which can create robustness–fragility tradeoffs in a population as a whole. Rather, assisted gene flow is a form of managed breeding that can be conducted immediately in coral populations to help conserve the overall genetic diversity of an entire species. The ability to use cryopreserved coral sperm in this process makes AGF even more feasible in the near term, and the ability to bank cryopreserved coral sperm indefinitely will make this an even more powerful option over the long term.

### Considerations for Future Outplanting of AGF Corals

Outplanting, the process of moving a cultured coral colony to a natural reef for restoration purposes, is the ultimate goal of many coral cultivation efforts. Permitting protocols for outplanting are generally well-established for *in situ* nursery-reared corals, and these typically require compliance with best practice guidelines and adherence to a strict monitoring and reporting schedule (Lirman and Schopmeyer 2016). However, the outplanting of *ex situ*-reared colonies presents a novel set of considerations for permitting, particularly when colonies are reared or moved for AGF purposes, and the permitting requirements for such projects are already under consideration by some policy makers (Great Barrier Reef Marine Park Authority 2018). In general, when developing and issuing permits for AGF projects, the risk of negative genetic consequences such as outbreeding depression can be considered and mitigated beforehand with thorough knowledge of population structure and data on population-level inbreeding coefficients (Aitken and Whitlock 2013). Furthermore, veterinary examinations can be used to minimize disease risks associated with introduced organisms (Viggers et al. 1993). Health examinations are often prerequisite for the release of any marine organism maintained in an *ex situ* system (Florida Fish and Wildlife Conservation Commission 2009), and veterinary protocols have been established for the introduction of *in situ*- and *ex situ*-reared corals to natural reefs (Berzins et al. 2008). Equivalent protocols could be developed and expanded to encompass the specific issues related to corals reared using AGF, including the use of colonies that have been selectively bred to possess desirable phenotypic traits (van Oppen et al. 2017). Additionally, broad best practice recommendations for conservation programs using AGF are already available for terrestrial species (O’Neill et al. 2008), and these could serve as a framework for the future development of best practices guidelines for coral AGF.

## Conclusion

This study provides the first evidence that gene flow is possible between genetically-isolated populations of the threatened Caribbean coral *A. palmata*, and in doing so, demonstrates that cryopreserved sperm can be used to move genetic diversity in this species. This proof-of-concept work represents an important step forward for coral reproductive biology, both regionally and worldwide, by demonstrating the feasibility of cryopreservation-assisted gene flow. Further, this study creates the foundation for future experiments in which sperm from thermally-tolerant corals could be used to transfer heritable thermal tolerance to other coral populations. The complexity of carrying out these experiments over time and space highlights the importance of capacity-building across the Caribbean region so that future cryopreservation-based experiments and restoration efforts can be carried out with more species, in new locations, and at greater scales.

## Acknowledgements

We are grateful to the principal funders behind this project, the Paul G. Allen Philanthropies, for their vision and willingness to take a risk on an ambitious, first-time project. We received valuable logistical support and advice from colleagues at NOAA, CARMABI, Smithsonian Institution, Mote Marine Laboratory, and the U.S. Fish and Wildlife Service office in Miami. S. Graves and D. Vaughan helped procure funding and provide space for the project. M. Miller provided valuable advice on the feasibility and rationale for the project. C. Wilson at USDA National Animal Germplasm Program helped with the storage and shipment of cryopreserved samples. A. Moura and S. Winters at Coral Restoration Foundation helped with shipping logistics and Airgas Miami kindly contributed liquid nitrogen fills during transit. In Curaçao, we are grateful for the support, field assistance, and assistance in permitting, diving, and shipping that we graciously received from F. Dilrosun at GMN, C. Winterdaal at CARMABI, the Baums Lab, and the local staff of the Diveshop Curaçao, Curaçao Sea Aquarium, Swissport, American Airlines, FedEx, Curaçao Customs, and Lindegas N.V. Student interns and volunteers at CARMABI assisted with large-scale dive preparations and gamely accommodated the ambitious scale of the project. We are especially thankful to the divers who participated in anywhere between one and thirty *Acropora* monitoring and collection dives at CARMABI in 2018, including R. C. Barnes, I. Baums, G. Boecker, K. Fitzgerald, R.-J. Geertsma, S. Gordon, M. Hooftijzer, E. Houtepen, J. Huckeba, K. Latijnhouwers, K. Leeper, A. Musk Schantz, K. Stankiewicz, C. van Bijnen, and K. Vasquez Kuntz. To obtain the sperm samples from genetically-distinct populations of *A. palmata*, SECORE divers and volunteers helped collect *A. palmata* samples from Puerto Rico in 2008, and divers and volunteers for CRF and The Florida Aquarium Center for Conservation helped collect *A. palmata* samples from Florida in 2016. These were processed and cryopreserved by members of the Hagedorn Laboratory and members of the Coral Cryopreservation Master Class. All cryopreserved samples were shipped to the USDA National Animal Germplasm Program, to whom we are grateful for assistance with long-term sample storage. Cryopreserved samples were exported to Curaçao under CITES Certificate #18US59512C/9. Coral larvae were exported to Florida under CITES permits #18CW007 and #18CW012 issued by the Curaçao Department of Health, Environment, and Nature (Curaçao Ministerie van Gezondheid, Milieu, en Natuur). J. Bouwmeester assisted with data compilation. In addition to the support from the Paul G. Allen Family Foundation, MH was further supported in her work by the Smithsonian Conservation Biology Institute, the Hawaii Institute of Marine Biology, the Paul M. Angell Family Foundation, the Roddenberry Foundation, the Seaver Institute, the William H. Donner Foundation, the Barrett Family Foundation, the Skippy Frank Foundation, the Compton Foundation, the Cedar Hill Foundation, and the Anela Kolohe Foundation. The Volgenau-Fitzgerald Family Fund supported the creation of a science education film about this project. KLM, MJAV, and the dive and lab teams at CARMABI were further supported in the field through grants from National Geographic and private donations, as well as by administrative support from Multiplier Project Accelerator, and by annual subsidies provided to CARMABI by the Government of Curaçao. Project funders did not influence the interpretation of the project data or the decision to publicize the findings. Finally, we are grateful to populations of *Acropora palmata* around the Caribbean, which contributed over one hundred thousand eggs and tens of billions of sperm cells to this project.

## Author Contributions

MH, KLM, KO, CAP, TV, JM, and TM designed the project and acquired the funding. MH, CL, and NZ designed and optimized the cryopreservation methods and tools that were used in the experiments. MH and CL led all cryopreservation steps and trained KLM, LT, DMF, MJAV, and VFC in cryopreservation methods on site at CARMABI. KLM and VFC led all field logistics at CARMABI including spawning dives, lab preparations, materials preparation, and training in larval husbandry with help from MJAV, LT, DMF, MH, and CL. KLM organized the cryopreservation logistics at CARMABI and designed the procedures for larval shipping. KLM, DMF, LT, VFC, and MJAV dove and collected the gamete bundles from *Acropora palmata*. MH, KL, and CL oversaw the cryopreservation of samples in Curaçao and the thawing and post-thaw analyses of samples. DMF, LT, KLM, and MJAV prepared eggs for fertilization, set up the *in vitro* fertilization experiments, cleaned the embryos, reared the larvae, and prepared the larvae for shipping. KO and CAP prepared aquarium systems and settlement substrates, received the larvae, settled the larvae, and performed post-settlement husbandry, care, grow-out, and monitoring. KLM, MH, DMF, LT, KL, and CL collected, compiled, and analyzed data on gamete production, sperm motility, sperm concentration, freezing rates, post-thaw motility, and fertilization rate. KO, TM, JM, KLM, MH, and MJAV obtained the transport permits and coordinated logistics for customs inspections and clearances. KLM and MJAV documented the project activities for outreach purposes. MH, KLM, KL, CAP, and KO wrote the paper with input from MJAV, DMF, LT, and VFC.

